# The origins of anterograde interference in visuomotor adaptation

**DOI:** 10.1101/593996

**Authors:** Gonzalo Lerner, Scott Albert, Pedro A. Caffaro, Jorge I. Villalta, Florencia Jacobacci, Reza Shadmehr, Valeria Della-Maggiore

## Abstract

Anterograde interference refers to the negative impact of prior learning on the propensity for future learning. Previous work has shown that subsequent adaptation to two perturbations of opposing sign, *A* and *B*, impairs performance in *B*. Here, we aimed to unveil the mechanism at the basis of anterograde interference by tracking its impact as a function of time through a 24h period. We found that the memory of *A* biased performance in *B* for all time intervals. Conversely, learning from error was hindered up to 1h following acquisition of *A*, with release from interference occurring at 6h. These findings suggest that poor performance induced by prior learning is driven by two distinct mechanisms: a long-lasting bias that acts as a prior and hinders the initial level of performance, and a short-lasting learning impairment that originates from a reduction in error-sensitivity. Our work provides insight into the timeline of memory stabilization in visuomotor adaptation.

## INTRODUCTION

We gain robustness through adaptation: in the face of environmental and/or internal perturbations, adaptation maintains the precise control of elementary movements like reaching and saccades. Like other types of learning, adaptation may lead to interference or facilitation depending on the level of congruency of sequentially learned materials. Facilitation of learning is commonly referred to as savings, a process by which subsequent exposure to the same perturbation results in faster learning (Krakauer, 2009). In contrast, successive adaptation to opposing perturbations (e.g., rotation *A* followed by rotation *B*) may lead to a deficit in the learning of *B*. This phenomenon, known as anterograde interference, has been reported in visuomotor and force-field adaptation paradigms when successively adapting to conflicting perturbations within the same reaching task (Brashers-Krug, Shadmehr, & Bizzi, 1996; Sing & Smith, 2010; Tong & Flanagan, 2003; Wigmore, Tong, & Flanagan, 2002). Yet, there is currently no consensus on whether anterograde interference is transient or long lasting. In fact, whereas some studies suggest that anterograde effects may last less than a few hours (e.g., Brashers-Krug et al., 1996; Thoroughman & Shadmehr, 1999), others appear to point to a long lasting impact in the time scale of days (Caithness et al., 2004; Miall, Jenkinson, & Kulkarni, 2004). It has even been suggested that anterograde interference may be stronger than retrograde interference (Caithness et al., 2004; Miall et al., 2004; Sing & Smith, 2010), masking the effect of interest in retrograde protocols aimed at unveiling the time course of memory consolidation (Miall et al., 2004).

This lack of consensus may be partly due to the manner in which anterograde interference is quantified (Sing, Joiner, Nanayakkara, Brayanov, & Smith, 2009). Previous studies estimated the amount of interference predominantly based on the initial level of performance, computed by averaging through the first trials of the learning curve. This empirical measure does not discriminate between changes in learning rate and retention. That is, initial performance in *B* is a mixture of how much the subject has retained what they learned in *A*, and how much they can learn from errors experienced in *B*. If anterograde interference arises from impairment in the ability to learn, one would expect that prior exposure to *A* would reduce the learning rate in *B*. Yet, with the exception of Sing & Smith (2010) no study that we are aware of has focused on the rate of learning as the fundamental measure of anterograde interference.

Here, we aimed to unveil the origins of anterograde interference by varying the time interval elapsed between adaptation to opposing rotations through a 24 h period. Our experimental approach allowed us to estimate the contribution of a prior memory independently from the ability to learn. The recruitment of a large number of subjects (n = 93) allowed us to analyze and contrast individual measures of learning during adaptation to *A* and *B*, when the two events were separated by 5 min, 1 h, 6 h and 24 h. In addition, we analyzed cycle-by-cycle learning using a state-space model (Albert & Shadmehr, 2018; Cheng & Sabes, 2006; Donchin, Francis, & Shadmehr, 2003; Ethier, Zee, & Shadmehr, 2008; Smith, Ghazizadeh, & Shadmehr, 2006). This allowed us to identify the impact of anterograde interference on three variables: initial state, error-sensitivity, and retention.

We found that poor performance observed when *A* and *B* are learned successively is driven by two distinct phenomena operating on different time scales: the influence of a prior memory that biases initial behavior, and an impairment in the ability to learn from errors in the new context. Whereas the former appears to impact behavior on a scale of days, the latter resolves within a 6 h period.

## MATERIALS AND METHODS

### Participants

Ninety-three healthy volunteers (33 males; ages: mean ± std. dev. 24±4 years old) with no known history of neurological or psychiatric disorders were recruited from the School of Medicine of the University of Buenos Aires. All subjects were right handed as assessed by the Edinburgh handedness inventory (Oldfield, 1971). The experimental procedure was approved by the local Ethics Committee and carried out according to the Declaration of Helsinki.

### Experimental Paradigm

Subjects were seated in a comfortable chair and performed a center-out-back task using a joystick operated with the thumb and index fingers of their right hand. Visual information was presented on a computer screen. The right elbow laid comfortably on an armrest and the wrist laid on a structure that fixed the joystick over a desktop. Subjects were told to maintain the same wrist posture across experimental sessions. Vision of the hand was occluded throughout the study.

At the beginning of each trial, we displayed one of 8 potential targets (0.4 cm diameter, placed 2 cm from the start point and concentrically located 45° from each other) on a computer screen. Joystick position was represented on the screen with a grey cursor of the same size as the target. The gain of the joystick was set to discourage subjects from correcting their movements online. Specifically, a displacement of 1.44 cm of the tip of the joystick moved the cursor on the screen by 2 cm. On average, movement time for correct trials was 125.5 ± 26.6ms (mean ± 1 std. dev.), providing little or no opportunity for within-movement corrections based on visual feedback. Participants were instructed to make a shooting movement through the target, as fast as possible, starting at target onset. There were 8 trials per cycle (one for each target) and 11 cycles per block. The order of target presentation was randomized within each cycle.

Two types of trials were presented throughout the experimental session (Fig. 1A). During *null* trials, participants performed shooting movements in the absence of a perturbation. During *perturbed* trials, a counterclockwise (CCW, labeled as perturbation *A*), or a clockwise (CW, labeled as perturbation *B*) visual rotation of 30° was applied to alter the trajectory of the cursor.

**Figure 1.**
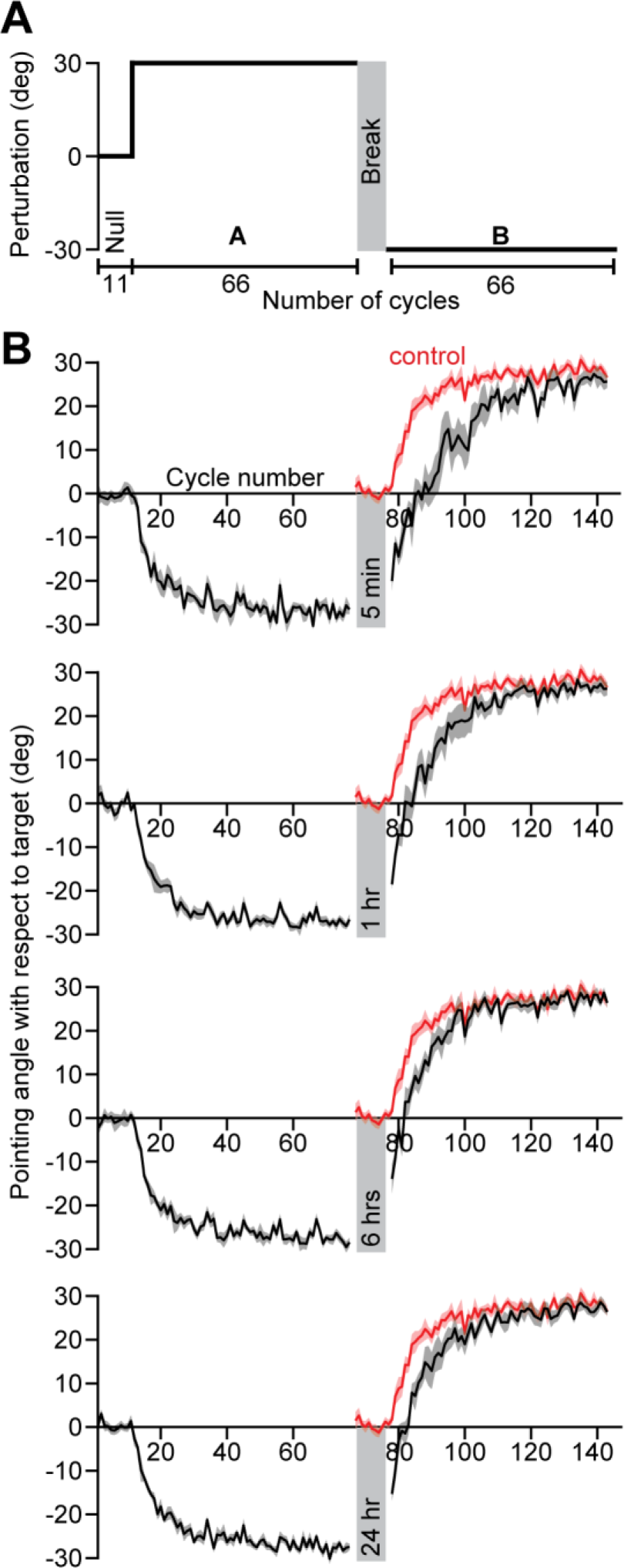
Experimental paradigm and learning curves. **A.** Paradigm. Subjects held a joystick and made pointing movements towards one of eight visual targets shown on a display. The experiment began with 11 cycles of null trials (Null) after which a 30° counterclockwise rotation was applied to the cursor for 66 cycles (*A*). Next, each experimental group waited a different length of time ranging from 5 min to 24 h. After this break, subjects were immediately exposed to a −30° clockwise rotation (*B*) for 66 cycles. **B.** Behavior. Subject pointing angles on each trial were collapsed into cycles by identifying the median pointing angle across each cycle of 8 trials. Each inset shows the median behavior of 1 of the 4 experimental groups. The shaded region indicates ±1 standard error of the median. Each group differs in the amount of time that elapsed between the exposure to the *A* and *B* periods (from top to bottom: 5 min, 1 h, 6 h, and 24 h). The behavior for each group (black) is compared with that of a control group (red) that was exposed to 11 cycles of null and then 66 cycles of the *B* rotation.

Feedback about the subject’s movement was provided on each trial via the color of the cursor, which varied along a gradient between red (miss) and green (hit). Furthermore, subjects had a limited amount of time to complete the movement after the appearance of the target. If the elapsed time exceeded 900 ms, the trial was aborted and the cursor was turned red until the next trial. Target hits with error < 2.5° were rewarded with a simulated sound of an animated explosion. The total score (hit percentage) was displayed on the screen at the end of each block. Subjects were instructed to try to maximize this score throughout the experiment. The task was programmed using MATLAB’s Psychophysics Toolbox, Version 3 (Brainard, 1997).

### Experimental Procedure

Figure 1A illustrates the experimental design. Participants were randomly assigned to one of four experimental groups or a control group. The experimental groups (Fig. 1A) performed one block (11 cycles) of null trials followed by six blocks (66 cycles) of CCW perturbed trials (perturbation *A*). After a variable time interval, they performed six blocks (66 cycles) of the CW perturbation (perturbation *B*). The four experimental groups were distinguished by the amount of time that separated the two rotations: 5 min (n = 16), 1 h (n = 20), 6 h (n = 19), and 24 h (n = 18). This variation in the period between perturbations *A* and *B* allowed us to assess how the passage of time impacted on the initial level of performance in *B* (first cycle), as well as on each subject’s ability to adapt to *B*.

A group of subjects (n = 20) experienced only the *B* perturbation. This control group served two purposes. First, it was critical for our analysis of anterograde interference, serving as our benchmark for performance in *B* without any potential influence of learning in *A*. Second, given that subjects always learned *A* before *B*, this group was key in ruling out an order effect. Control subjects performed 1 block (11 cycles) of null trials followed by 6 blocks (66 cycles) of *B*.

### Data post-processing

For each trial, the pointing angle was computed based on the angle of motion of the joystick relative to the line segment connecting the start and target positions. Trials in which pointing angles exceeded 120° or deviated by more than 45° from the median of the trials for each cycle were excluded from further analysis (1.6% of all trials). After this processing, the trial-by-trial data were converted to cycle-by-cycle time series by calculating the median pointing angle in each 8-trial cycle for each subject. Unless otherwise noted, the cycle-by-cycle data were used for each analysis reported in this work.

### Model-free data analysis

We empirically quantified each subject’s learning rate in *A* and *B* by fitting a single exponential function (Eq. 1) to the pointing angle corresponding to the *A* and *B* periods.

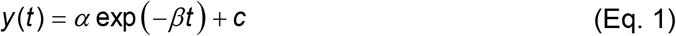

Here *y*(*t*) represents the pointing angle on cycle *t*. The first cycle of the rotation was represented by *t* = 0. The exponential fits included three parameters. The parameters *α* and *c* determine the initial bias and the asymptote of the exponential, respectively. The parameter *β* represents the learning rate of the subject. We constrained the relationship between *α* and *c* to force the exponential fit to intersect subject behavior at time step t = 0. Therefore, the exponential function had only two free parameters; the third was fixed by the initial level of subject performance. We fit one exponential function to the 66 cycles of the *A* rotation and another one to the 66 cycles of the *B* rotation (Figure 1). Each period was fit using the *fmincon* function (MATLAB 2018a) to minimize the squared error between subject behavior and the exponential fit.

Although the exponential function closely approximates the decay of motor error during adaptation to a single perturbation, its learning rate parameter reflects a mixture of cycle-by-cycle forgetting and error-based learning. This potentially confounds our analysis of interference because during the *A* perturbation, the direction of forgetting (always towards baseline performance) opposes the direction of error-based learning. However, during the *B* perturbation, an initial bias in the performance of the experimental groups towards *A* causes forgetting and error-based learning to act in the same direction. This relationship switches once subjects pass the “zero point” of baseline performance: here retention and error-based learning again oppose one another. These considerations illustrate the difficulties inherent in using exponential fits to disambiguate the differential effects learning and forgetting may have on behavior.

### State-Space Model

To better quantify subject performance in *A* and *B*, we used a state-space model that dissociates the effect of cycle-to-cycle learning from forgetting while appreciating initial biases in learning.

When people perform a movement that produces an unexpected result, they learn from their movement error and retain part of this learning over time. In other words, behavior during sensorimotor adaptation can be described as a process of error-based learning and trial-by-trial forgetting (Donchin et al., 2003; Smith et al., 2006; Thoroughman & Shadmehr, 2000). State-space models of learning consider how the behavior of a learner changes due to trial-by-trial error-based learning, and decay of memory due to the passage of time (i.e., trials). To examine the anterograde interference of *A* on *B*, we fit a single module state-space model to the empirical data. This allowed us to ascribe any differences in performance during the *B* period to meaningful quantities: sensitivity to error, forgetting rate, and initial state.

We imagined that the state of the learner (the internal estimate of the visuomotor rotation) changed from one cycle to the next, due to error-based learning and partial forgetting, according to Eq. (2).

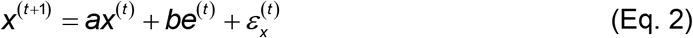

Here *x*^(*t*)^ represents the state of the learner on cycle *t*. The parameter *a* is a retention factor that encapsulates how well the subject retains a memory of the perturbation from one cycle to the next. The parameter *b* represents sensitivity to error and determines the rate at which each subject learns from error. The error sensitivity is multiplied by the visual error *e*^(*t*)^ between the pointing angle and target. The change in state from one cycle to the next is corrupted by state noise 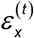 which we assumed to be Gaussian with mean zero and variance equal to 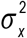.

The internal state of the subject is not a measurable quantity. Rather, on each cycle, the motor output of the subject is measured. We imagine that the motor output directly reflects the internal state but is corrupted by motor execution noise according to Eq. (3).

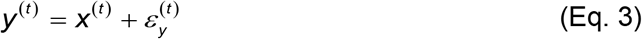

As with our exponential fit of Eq. (1), here *y*^(*t*)^ represents the subject’s pointing angle on cycle *t*. We assumed that the motor execution noise 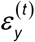 corrupting the reaching movement was Gaussian with mean zero and variance equal to 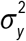.

We fit the state-space model to cycle-by-cycle single subject behavior using the Expectation Maximization (EM) algorithm (Albert & Shadmehr, 2018). The algorithm identified the parameter set that maximized the likelihood of observing each sequence of subject pointing angles given the parameters and structure of our state-space model. This parameter set contained 6 parameters: the retention factor *a*, error sensitivity *b*, state noise variance 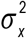, motor noise variance 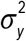, and two parameters describing the initial state of the learner. We modeled the initial state of the learner as a normally distributed random variable with mean *x*_1_ and variance 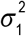. The parameter *x*_1_ served as our estimate of the initial bias of the learner.

To fit the model, we started the EM algorithm from 5 different initial parameter sets, performed 100 iterations of the algorithm (Albert & Shadmehr, 2018), and selected the parameter set with the greatest likelihood. We fit our state-space model to single subject behavior separately for the *A* and *B* periods. For the *A* period, we fit the 77 cycles encompassing the first 11 null cycles and the following 66 CCW rotation cycles (Fig. 1). We fit the initial null trials along with the perturbation trials to increase confidence in the model parameters. For the *B* period, we fit the 66 cycles encompassing the CW rotation (Fig. 1).

#### Validation of the single state-space model

Our primary analysis assumed that learning could be represented using a single adaptive state. For a single state system, impairment in the learning rate in *B* requires that the learning system (*i.e.*, the model parameters) has changed from the *A* to the *B* period. Prior accounts of anterograde interference considered how an impairment of learning might arise from the emergent properties of a two-state system (Sing & Smith, 2010). A two-state system might show a change in learning rate during the *B* period even if the system has not changed, simply due to differing initial biases in the underlying adaptive states of the system.

To validate the choice of a single state model, in a set of mathematical control studies, we tested if our findings were also consistent with the predictions of a two-state framework. For this analysis, we fit a two-state model of learning to the *A* and *B* sequences of subject pointing angles. The two-state model of learning is equivalent to the single state model of learning, with the exception that learning is described as the combined output of two parallel adaptive states, a fast learning state and a slow learning state. The states evolve over trials according to Eq. (4).

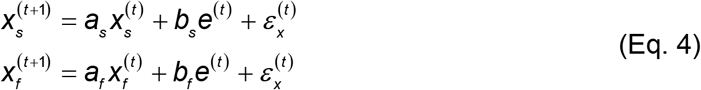

Here, the slow and the fast states are represented by the quantities *x*_*s*_ and *x*_*f*_, respectively. As with the single state model (Eq. 2) each state changes due to forgetting (described by its retention factor *a*) and error-based learning (described by its error sensitivity *b*). These internal estimates of the perturbation are additively combined to determine motor behavior according to Eq. 5.

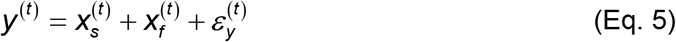

We fit this two-state model of learning to subject behavior during the *A* and *B* periods using the EM algorithm (Albert & Shadmehr, 2018). The algorithm identified the parameter set that maximized the likelihood of observing each sequence of subject pointing angles. We fit the model to the same cycles in *A* and *B* described for the single state model fits. To fit the model, we started the EM algorithm from 20 different initial parameter sets, performed 250 iterations of the algorithm, and selected the parameter set with the greatest likelihood. The model parameter set consisted of 9 variables: slow and fast retention factors *a*_*s*_ and *a*_*f*_, slow and fast error sensitivities *b*_*s*_ and *b*_*f*_, the variances of state evolution and motor execution, and three parameters for the initial state of the learner. We modeled the initial fast and slow states as normally distributed random variables with mean 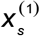 and 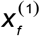, and variance 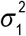. Each model was fit under the linear constraints *b*_*f*_ > *b*_*s*_ and *a*_*s*_ > *a*_*f*_. These constraints enforce that the slow state learns more slowly than the fast state, but also retains its memory better from one trial to the next (Smith et al., 2006).

We compared the single state model and two-state model in their abilities to describe subject behavior. To compare these models, we computed the Bayesian Information Criterion (BIC) according to Eq. 6.

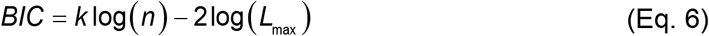

Here *k* represents the number of model parameters (6 for the single state model, 9 for the two-state model), *n* represents the number of data points, and *L*_max_ represents the maximum likelihood for the model fit obtained using the EM algorithm. To obtain a single estimate of *BIC* for each subject, we averaged the BIC over the *A* and *B* periods. To quantify the evidence for each model, we compared the BIC distributions for the single state and two-state models for all subjects in the experimental groups using a paired *t*-test.

Finally, to test the hypothesis (Sing & Smith, 2010) that a learning impairment in *B* could trivially arise from the interplay between fast and slow state dynamics (rather than an actual impairment in the learning system) we simulated behavior during the *B* period using a two-state model, and compared this behavior to actual subject behavior. We reasoned that if the learning system did not differ across the *A* and *B* periods, the *B* period behavior should be predicted by the model parameters fit to the *A* period. To test this idea, we used the model parameters fit to the *A* period to simulate individual subject behavior during the *B* period, focusing specifically on the 5 min group that demonstrated the greatest amount of interference.

To simulate behavior in the *B* period, we calculated the expected value of subject performance by removing the noise terms in *Eqs*. 4 and 5. Due to the passage of time, behavior at the start of the *B* period exhibited decay relative to the performance at the end of the *A* period. We accounted for this forgetting using the initial biases in the fast and slow states. To do this, we calculated the amount of forgetting that occurred from the last cycle in *A* to the first cycle in *B*, as a percentage. We then estimated the final levels of the slow and fast states in *A* by simulating behavior in *A* using *Eqs.* 4 and 5, without any noise terms. For our simulation of the *B* period, we set the initial fast and slow state levels to the final levels in *A*, scaled down by the forgetting percentage.

We calculated the rate of learning using our exponential model (*Eq.* 1) of behavior both for the actual behavior recorded in *B* and the behavior simulated using the two-state model. We compared these rates using a paired *t*-test to determine how well the two-state parameters from the *A* period accounted for the interference observed in the *B* period.

### Statistical assessment

Statistical differences were assessed at the 95% level of confidence. Prior to statistical testing, outlying parameter values were detected and removed based on a threshold of three median absolute deviations from the group median. For cases where our variables of interest did not fail tests for normality and equality of variance, we used a one-way ANOVA for our statistical testing. In cases where the statistical distributions failed tests for both equal variance across groups (Bartlett’s test) and normality (Shapiro-Wilk test) we used the Kruskal-Wallis test to detect non-parametric differences across experimental groups. In cases where our statistical tests indicated a significant effect of group (p < 0.05), we used either Tukey’s test (following one-way ANOVA) or Dunn’s test (following Kruskal-Wallis) for post-hoc testing. For the latter, pairwise tests of all experimental groups were conducted against the control group and Bonferroni corrected. In cases where one-way ANOVA was used for statistical testing, complementary figures depict the mean statistical quantity for each group as well as the standard error of the mean, calculated assuming a normal distribution. In cases where Kruskal-Wallis was used for statistical testing, complementary figures depict the median statistical quantity for each group as well as the standard error of the median (estimated with bootstrapping). When comparing mean values against zero, a one-sample t-test test was used followed by the Bonferroni correction for multiple comparisons.

## RESULTS

When people adapt to perturbation *A*, and then switch to the opposite perturbation *B*, performance in *B* appears impaired (Brashers-Krug et al., 1996; Braun, Aertsen, Wolpert, & Mehring, 2009; Caithness et al., 2004; Shadmehr & Brashers-Krug, 1997; Tong & Flanagan, 2003). Pinpointing the origin of this behavioral deficit is difficult because performance in *B* may reflect two different processes: the level of retention of the memory of *A*, and the ability to learn *B*. In addition, these factors may vary independently as a function of time. Our study aimed to dissociate between these two factors by varying the time interval elapsed between *A* and *B* as subjects adapted to conflicting visuomotor rotations.

On each trial, subjects moved a joystick to displace a cursor to one of 8 targets. On average, movement time for correct trials was 125.5 ± 26.6ms (mean ± 1 std. dev.), providing little or no opportunity for within-movement corrections. All groups initially trained in a baseline period of null trials (no perturbation) followed by adaptation to perturbation *A* (Figure 1A). After completion of training in *A*, subjects waited for a specific amount of time (5 min, 1 h, 6 h or 24 h), and then were exposed to perturbation *B*. Figure 1B shows the pointing angle during null trials (cycles 1 to 11), learning of *A* (cycles 12 to 77) and learning of *B* (cycles 78 to 143) for each of the experimental groups (black curves) and the control group (red curve). Pointing angle refers to angle of motion of the joystick relative to the line segment connecting the start and target positions. As expected, the pointing angle during null trials was close to zero. During exposure to perturbation *A*, subjects shifted their pointing angle gradually, asymptotically approaching −30° (Shmuelof et al., 2015). After adapting to *A* and waiting the assigned time, subjects returned and were exposed to perturbation *B*.

How did learning of *A* impact performance in *B*? We quantified the initial level of performance in *B* as the mean pointing angle during the first cycle of adaptation for each group (Fig. 2A). Given that little or no learning is expected to take place in one cycle (1 cycle = one trial per target), this measure allowed us to estimate the recall of *A*. The initial level of performance in *B* was biased towards *A*, and decayed as a function of time (Fig. 2A, one-way ANOVA, F_(69,3)_ = 3.37, p = 0.029; Tukey’s test, 5 min marginally different from 6 h with p = 0.073, 5 min different from 24 h with p = 0.029, all other comparisons have p > 0.358). Notably, even at 24 h the memory of *A* remained strong, exhibiting nearly 50% retention (one-sample t-test against zero with Bonferroni correction: p < 0.001 for all groups). This observation is consistent with the presence of a lingering memory of *A* (Shadmehr & Brashers-Krug, 1997; Thoroughman & Shadmehr, 1999).

**Figure 2.**
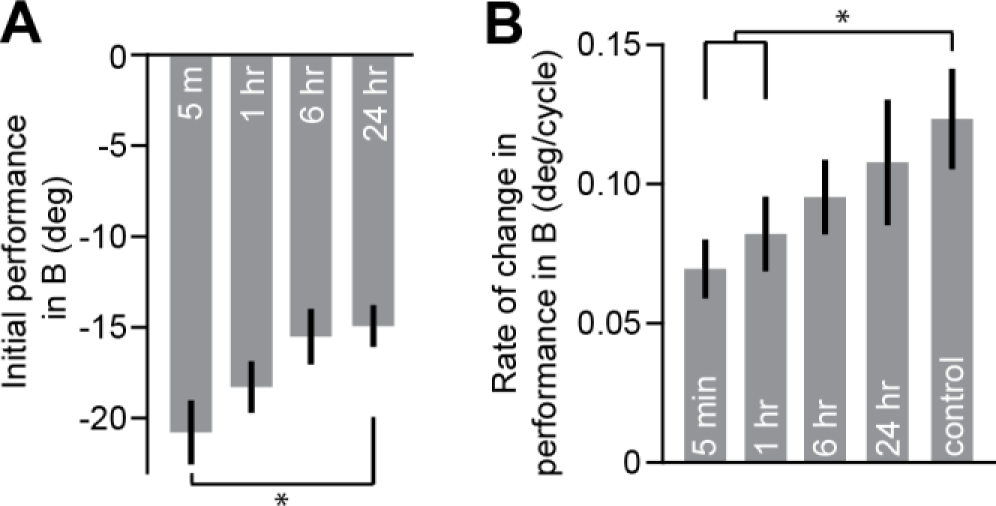
Effect of prior learning on different behavioral parameters. **A.** The initial level of performance in *B*, estimated from the mean pointing angle on the first cycle is displayed here for all groups. Given that learning within one cycle is minimal, this measure reflects the retention of the memory of *A*. **B.** The median rate of improvement (*i.e.*, the empirical learning rate) in *B* for all experimental groups and the control group is shown. In both **A** and **B**, experimental groups are shown from left to right in order of increasing temporal separation between the *A* and *B* periods. The control group is shown at far right where appropriate. In **A** and **B**, error bars indicate ±1 standard error of the mean and median, respectively. Asterisks indicate a level of significance of p < 0.05.

In summary, during initial performance in *B* the movements were strongly influenced by the presence of a memory of *A*. This memory decayed with time, but was still present at 24 h.

### Model-free analysis

In order to assess if having learned *A* also impaired the ability to learn *B*, we fit the motor output for each subject and each group during adaptation to *A* and *B* with an exponential function (Eq. 1). We found that during the *A* period there was no difference in the learning rates across the four experimental groups (Kruskal-Wallis, X^2^(62) = 4.75, p = 0.19). Therefore, the various groups were indistinguishable during learning of *A*.

Did prior exposure to *A* interfere with the rate at which *B* was acquired? To quantify the ability to learn *B*, we statistically compared the rate of change in performance for each experimental group during adaptation to *B* with that of the control group (Figure 2B). Non-parametric testing revealed a significant effect of group on the ability to learn *B* (Fig. 2B; Kruskal-Wallis, X^2^(80) = 10.84, p = 0.029). Post-hoc comparison between each experimental group and the control group identified a significant difference at 5 min and 1 h (Dunn’s test with Bonferroni correction, 5 min different from control with p = 0.044, 1 h different from control with p = 0.024), that disappeared by 6 h (6 h and 24 h not different from control with p > 0.952). This temporal pattern in the impairment of motor learning is consistent with anterograde interference (Krakauer, 2009).

To visualize differences in the learning of *B*, we artificially aligned the performance of the control group to each experimental group by shifting the control learning curve along the time axis (Figure 3). This procedure makes use of a fundamental property of exponential functions:

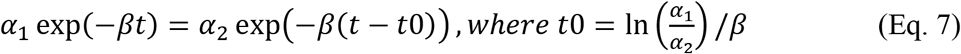

**Figure 3.**
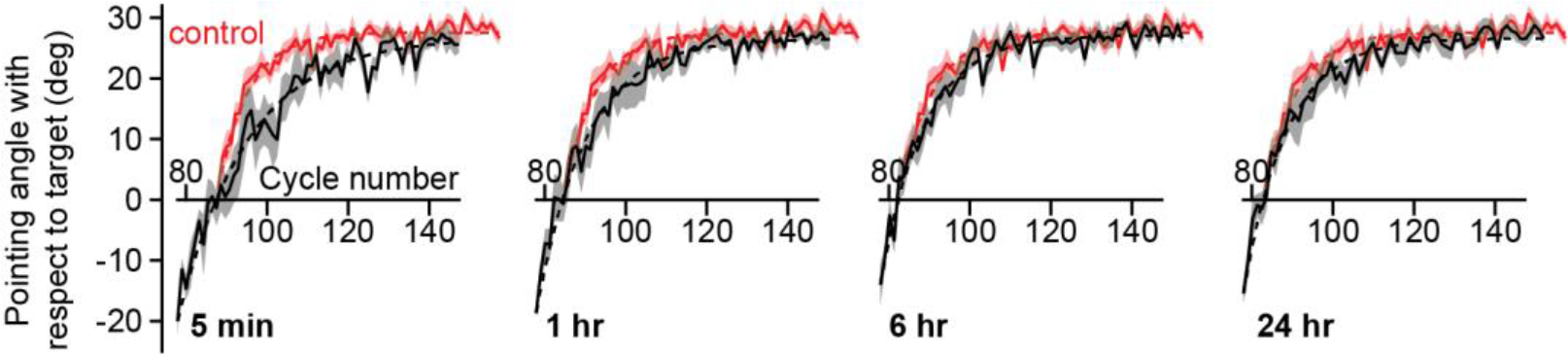
Anterograde interference and performance in *B*. The median pointing angle of each experimental group during learning the *B* rotation (solid black) and the control group (solid red) is shown. The shaded error region indicates ±1 standard error of the median. To highlight any difference in the rate of learning of the *B* rotation, the behavior of the control group was shifted in all plots, so that it roughly intersected the behavior of each experimental group at a pointing angle of approximately 0°. The magnitude of the shift was determined by fitting exponential curves (dashed lines) to the behavior of the experimental group (black) and control group (red) and horizontally shifting the control group so that the exponential fits intersected at 0°. From left to right, experimental groups are ordered in terms of increasing separation between the *A* and *B* perturbations.

If two exponentials start at different points (*α*_1_ *and α*_2_) but share the same decay rate *β*, then shifting one in time by *t*0 will result in complete overlap of the two functions.

To optimally align the behavior of the control with that of the experimental groups, we fit an exponential function (Eq. 1) to the median behavior of each group. We next calculated the cycle on which the predicted behavior would intersect a pointing angle of 0° for each group, and shifted the behavior of the control to match that of each experimental group at Y = 0 (Fig 3). This temporal displacement points towards an impairment in the learning rate of the 5 min and the 1 h groups.

Finally, to rule out the possibility that our results may be explained by an order effect (subjects always learned the CCW rotation before the CW rotation), we statistically compared the rate of learning of the control group with those of the experimental groups during learning in *A*. No differences were found between the learning rates of *A* and *B* control (Kruskal-Wallis, X^2^(78) = 5.53, p = 0.237). Therefore, the control condition rules out the possibility that our results are explained by the order in which the perturbations were learned.

In summary, we conclude that while the lingering memory of *A* caused the starting point of the learning process to be strongly biased in all experimental groups, the learning process itself was impaired at 5 min and 1 h only. Release from interference, determined as the time point at which the rate of learning resembled that of the control group, took place about 6 h post adaptation.

### State-space model

The exponential model we employed for our empirical analysis implicitly assumed that the rate of learning remained constant across trials. For the *B* period, this assumption is unlikely to be true because initially, learning from the errors induced by *B* is aided by forgetting of the memory of *A*. That is, as the *B* period starts, performance falls toward baseline, and the rate of this fall is due to two processes: forgetting of *A*, and learning from error in *B*. During this period, forgetting and learning act in the same direction. However, once the performance crosses baseline levels, the influence of memory decay on behavior is in the opposite direction to learning from error. State-space models of learning disentangle these processes of forgetting and learning. For this reason, we fit a state-space model to the data from individual subjects separately during the *A* and *B* periods (Eq. 2 and 3).

The state-space model assumes that learning is governed by two processes: a process that learns from error, and a process that retains a fraction of that memory from one trial to the next. The state-space model closely tracked the observed behavior (Fig. 4). To quantify the model’s goodness-of-fit, we computed the fraction of each subject’s behavioral variance accounted for by our model fit (R^2^). To measure this coefficient of determination, we computed the expected value of the behavior predicted by our stochastic model (Eqs. 2 and 3) and compared this model prediction to each individual subject’s data. We found that across subjects, our model accounted for approximately 81.4 ± 8.4% (mean ± 1 std. dev.) of the variance in subject behavior. We repeated this analysis at the group level, where noise in the process of learning (Eq. 2) and production of a movement (Eq. 3) is smoothed over subjects. For each group, we computed the median behavior (Fig. 4A, black curves for experimental groups, red curve for control), the median behavior predicted by our model (Fig. 4A, blue curves for experimental groups, green for control), and then the coefficient of determination for these two time-courses. At the group level, the model accounted for 96.0 to 98.2% of the variance in median subject behavior.

**Figure 4.**
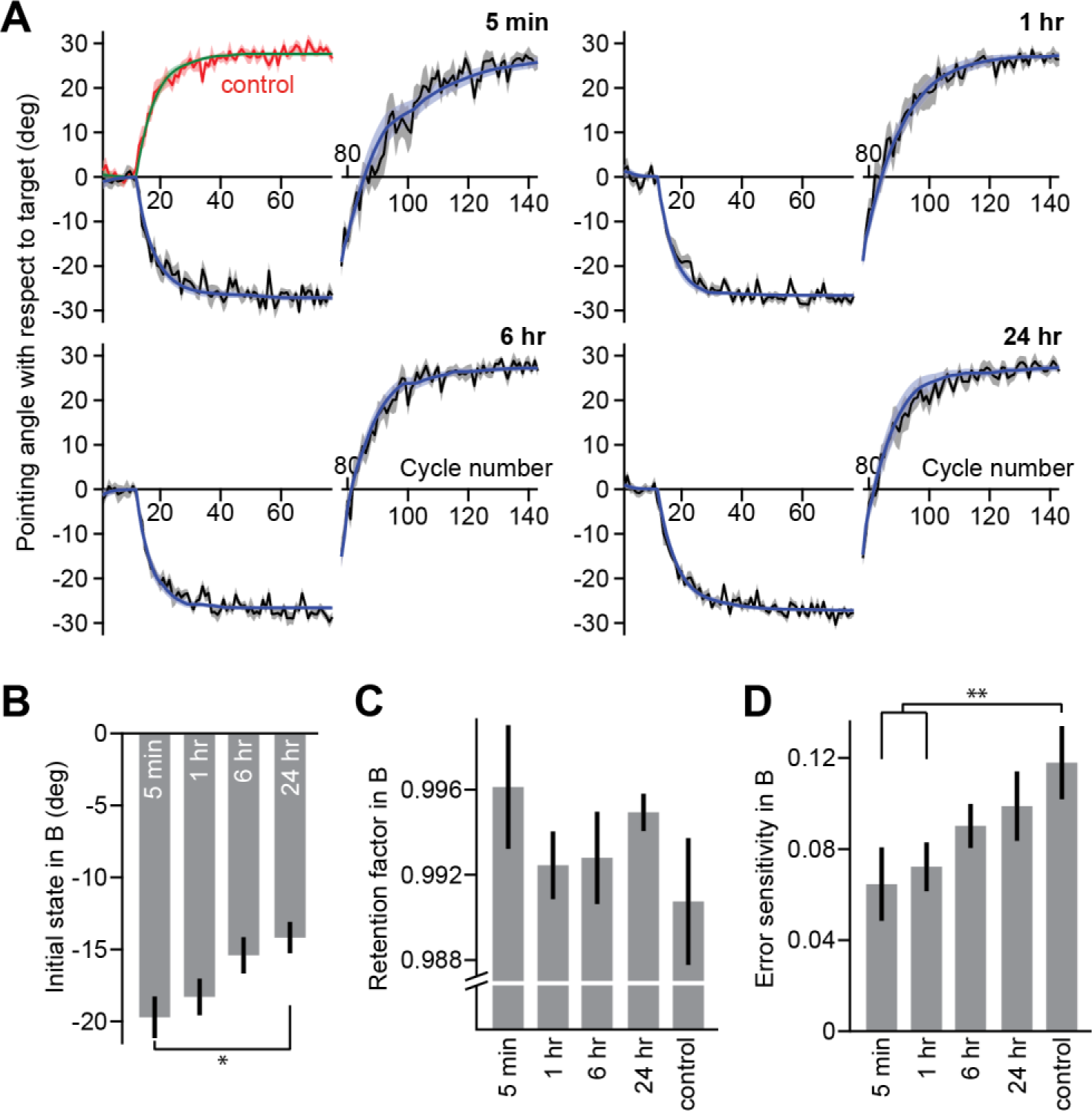
State-space model fit to behavior. **A**. We fit individual subject behavior using a single module state-space model of learning that accounted for cycle-by-cycle forgetting, error-based learning, and initial bias. We fit behavior separately for the *A* (cycles 1 through 77) and *B* (cycles 78 through 143) periods. Each plot depicts the median pointing angle for 1 of the 4 experimental groups (black lines) as well as the median pointing angle predicted from simulating the state-space model without noise (blue lines) using the maximum likelihood model parameter sets identified for each subject. Corresponding behavior (red) and state-space predictions (green) are provided for the control group in the top-left plot. The shaded error region indicates ±1 standard error of the median. **B**. Initial state. Here we report the initial state of the learner at the start of the *B* period. **C**. Here we report the retention factor during the *B* period for the experimental and control groups. **D**. We report the error sensitivity during the *B* period for the experimental and control groups. The height of each bar denotes the mean (**B**) or median (**C** and **D**) parameter value for each experimental group. Error bars indicate ±1 standard error of the mean or median. In all plots, experimental groups are shown from left to right in order of increasing temporal separation between the *A* and *B* periods. The control group is shown at far right where appropriate. Asterisks indicate a level of significance of p < 0.05 (*) or p < 0.01 (**).

After validating our model, we next considered how anterograde interference could be quantified at the level of three different processes that could affect performance in *B*: (1) memory of *A*, (2) cycle-to-cycle forgetting rate in *B*, and (3) learning from error in *B*. These three processes are represented separately by three specific model parameters: (1) the initial state of the learner in *B*, (2) the retention factor, and (3) the error-sensitivity.

Unsurprisingly, the initial state of the learner in *B* (Fig. 4B) closely followed our empirical estimate of the initial level of performance in *B* (Fig. 2A). As the interval between *A* and *B* increased, the initial state of the learner in *B*, *i.e.*, the amount of the *A* memory retained over time, decreased (one-way ANOVA, F_(69,3)_ = 3.89, p = 0.013; Tukey’s test, 5 min different than 24 h with p = 0.020, marginal differences between 5 min and 6 h as well as 1 h and 24 h with p = 0.097, all other comparisons have p > 0.347). However, despite this temporal decay, all groups retained at least 50% of the *A* memory (one-sample t-test with Bonferroni correction, all groups p < 0.001). Therefore, impairment of performance in *B* was in part caused by a lingering memory of *A* that did not fully decay even after 24 h.

To what extent was the impairment in *B* driven by changes in the rate of cycle-by-cycle memory retention and learning from error? Similar to our empirical analysis, we first confirmed that the experimental groups did not differ in performance during the *A* period. That is, there was no difference in the retention factor (Kruskal-Wallis, X^2^(66) = 0.53, p = 0.912) or error sensitivity (Kruskal-Wallis, X^2^(65) = 1.16, p = 0.763) across the experimental groups during adaptation to *A*. Furthermore, we found no difference in the retention factor during learning in *B* for any of the experimental groups, including the control group (Fig. 4C; Kruskal-Wallis, X^2^(79) = 5.66, p = 0.226). In contrast, error sensitivity was affected by prior learning in a manner consistent with anterograde interference (Fig. 4D; Kruskal-Wallis, X^2^(83) = 14.47, p = 0.006). Post-hoc tests against the control group unveiled a significant reduction in error sensitivity at 5 min and 1 h but not at longer time intervals (Dunn’s test with Bonferroni correction, 5 min different from control with p = 0.008, 1 h different from control with p = 0.004, 6 h and 24 h not different from control with p > 0.132).

In summary, our state-space model pointed to a similar conclusion drawn from our empirical findings. Prior exposure to *A* resulted in a bias in the initial state of *B* that persisted through 24 h. Moreover, prior exposure produced a reduction in error sensitivity, but this effect was short lasting: we found no evidence for it when the time between *A* and *B* was 6 h or more. Therefore, differences in performance in *B* for any timescale greater than 1 h were likely related to a prior memory of *A* and not to a deficit in learning.

### Validation of the computational approach

An earlier account of anterograde interference (Sing & Smith, 2010) investigated how the emergent properties of a learning system composed of two parallel adaptive states (Smith et al., 2006) could demonstrate impaired learning in *B* after the experience of *A*, even if there was no change in the learning rate of either state. Two-state models of learning posit that adaptation is supported by two parallel learning processes, a slow process (Fig. 5A, red) that learns little from error but exhibits strong retention over trials, and a fast process (Fig. 5A, green) that learns greatly from error but has poor ability to retain its memory over trials. Sing and Smith (2010), demonstrated that if a two-state learning system is exposed to the *A* perturbation (+30°), followed by the opposing *B* perturbation (−30°), learning of *B* would appear to be slowed (Fig. 5A, compare *B* curve with the naïve *A* curve shown in blue) because the slow state is heavily biased towards *A* at the start of *B*. That is, a two-state system can exhibit anterograde interference (a slowing of the overall adaptation rate), despite the fact that the individual learning rates have not changed from *A* to *B*. Could the reduction in error sensitivity we report in our analysis of anterograde interference be explained by a two-state system whose constitutive parameters do not differ across the *A* and *B* periods?

**Figure 5.**
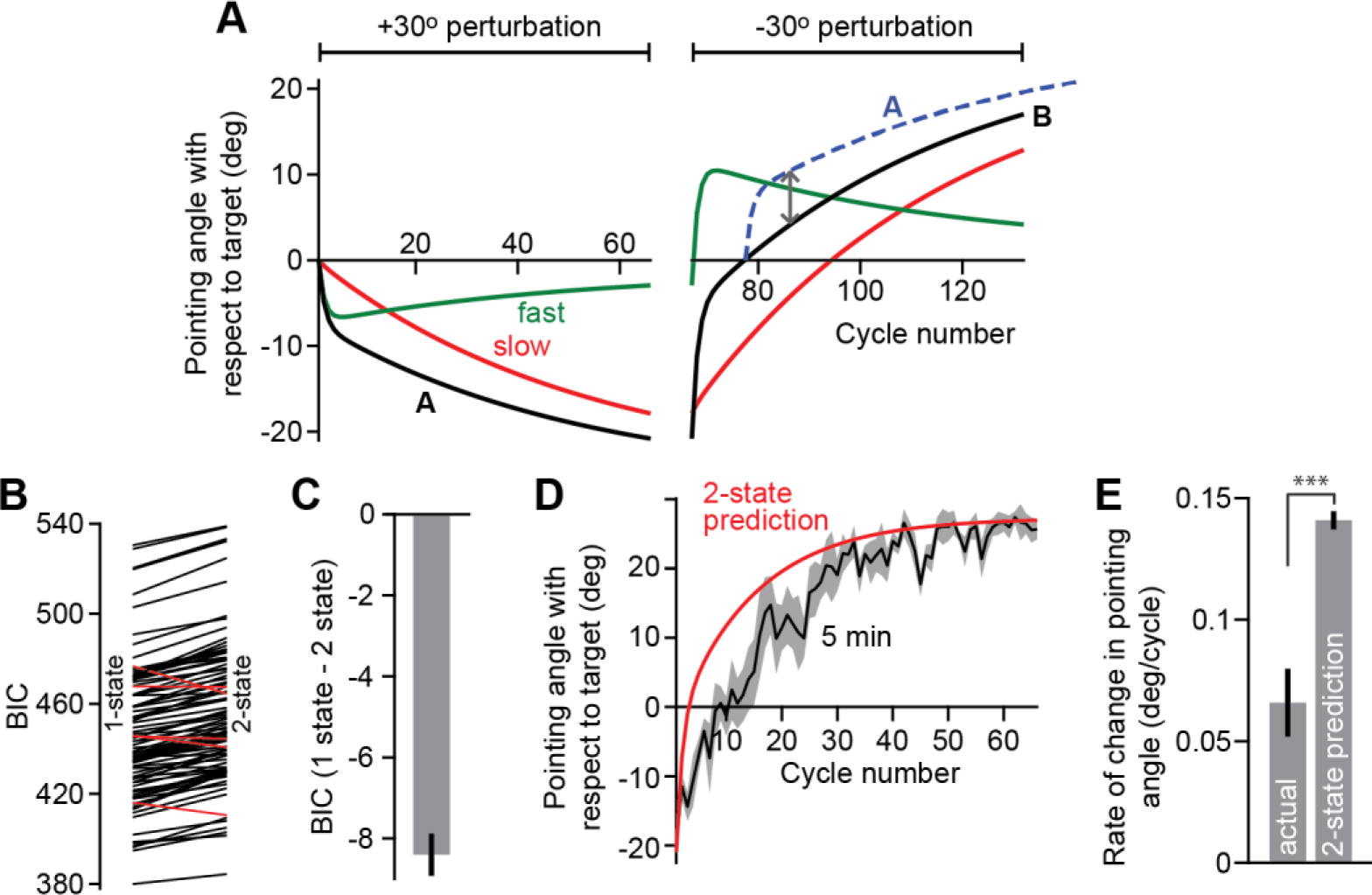
Validation of the state-space model. Prior accounts of anterograde interference have shown that a two-state model of learning can show an impairment in learning of the *B* memory without any change in the constitutive parameters of the two-state system from *A* to *B*. **A.** The two-state model (parameters obtained from Sing and Smith, 2010) posits that behavior (black) can be decomposed into parallel contributions from a slow (red) and fast (green) state. After learning the *A* perturbation (left half of the figure, +30° perturbation) both the fast and slow states are biased towards a memory of *A*. At the start of the *B* period (right half of the figure, −30° perturbation) the initial bias of the slow state towards *A* can slow the apparent rate of learning of the *B* perturbation, as compared to initial learning of *A* (blue trace shows the pointing angle during *A* with opposite sing and shifted to the 0° pointing angle of *B*). **B.** We asked if a two-state system could better account for subject behavior than the single state system considered in the primary analysis. We calculated the Bayesian Information Criterion (BIC) associated with single-state and two-state model fits to individual subject behavior. The endpoint of each line shows the average BIC for the *A* and *B* periods (left, single state model; right, two-state model). Each line depicts the result for a single subject. Red lines indicate subjects for which the two-state model was superior to the single state model. Black lines indicate subjects for which the single state model was superior to the two-state model. **C.** We calculated the difference in BIC for the single state and two-state models. Negative values indicate higher evidence for the single-state model. The height of the bar indicates the mean BIC, and error bars indicate ±1 standard error of the mean. **D.** We simulated behavior of each subject during the *B* perturbation (red) using the two-state model parameters fit to the *A* period. The initial bias of the slow state towards a memory of *A* did not produce learning in *B* that resembled the impaired rate of learning exhibited in the 5 min experimental group (black). Solid lines indicate the median prediction or behavior across subjects. Error shade indicates ±1 standard error of the median. **E.** To quantify any discrepancy between the rate of learning in *B* and the rate of learning predicted by the two-state model we fit our empirical exponential learning curve to the actual and simulated *B* behaviors. The height of each bar indicates the median empirical learning rate. Error bars indicate ±1 standard error of the median. Asterisks indicate a level of significance of p < 0.001.

To answer this question, we first mathematically compared the likelihood that our data was better described by a two-state system rather than a single state system. We fit a two-state model of learning (Albert & Shadmehr, 2018) separately for the *A* and *B* periods and compared the likelihood of this model with that of the single state model explored in our primary analysis using the Bayesian Information Criterion (BIC). At the level of individual subjects, we found that a two-state model of learning was justified in only 5 of the 73 subjects in the experimental groups (Fig. 5B, red lines). Therefore, in this task, the measured behavior was better described by a single state model of learning (Fig. 5C, lower BIC for single state model, paired t-test, t (92) = 16.133, p < 0.001).

We next asked if a two-state model fit to the *A* period of behavior would produce the pattern of interference we measured in *B*, as described by Sing and Smith (2010). For this analysis, we focused on the 5 min group that exhibited the most significant amount of anterograde interference (Fig. 2B). Using the two-state model parameters fit to the *A* period of behavior, we simulated subject performance during the *B* period. We found that the dynamics of learning predicted by a two-state model whose learning rates were unchanged from *A* to *B* (Fig. 5D, red), did not visually resemble the pattern of interference we measured in the 5 min experimental group (Fig. 5D, black). Indeed, the exponential rate of improvement in *B* exhibited by the two-state model simulation was a rather poor predictor of the actually measured behavior as exhibited by the 5 min group (Fig. 5E, paired t-test, t(15) = 4.235, p < 0.001). That is, our subjects learned much slower in the *B* perturbation than predicted by a two-state model in which parameters remain constant in *A* and *B*. This result gives us some assurance that the reduction in error sensitivity we observed during the *B* period was not due to a trivial property of a two-state learning system. Rather, it appears that following adaptation to *A*, the motor system was fundamentally impaired in its ability to learn *B*.

## DISCUSSION

How do motor memories influence one another? In this work, we studied the expression of anterograde interference in visuomotor adaptation by varying the time elapsed between learning opposing perturbations. We examined the impact of prior learning on the initial level of performance as well as the rate of learning over the time course of 5 min through 24 h. We found that these two parameters behaved very differently as a function of time. On one hand, adaptation in *A* biased the initial level of performance in *B*. Although the magnitude of this effect decreased with time, it remained strong at 24 h. On the other hand, prior adaptation hindered the ability to learn from error when perturbations were separated by 5 min and 1 h but not at 6 h and beyond, suggesting that actual learning is impaired on a shorter time scale, subsiding within a 6 h window.

There has been no general agreement in the literature of sensorimotor adaptation regarding how to define and, therefore, quantify anterograde interference. With the exception of Sing and Smith (2010), who measured the relative change in learning rate, most previous studies estimated anterograde interference based on the initial level of performance, by averaging across the first trials/cycles/blocks (Brashers-Krug et al., 1996; Krakauer, Ghez, & Ghilardi, 2005; Lee & Schweighofer, 2009; Shadmehr & Brashers-Krug, 1997; Tong & Flanagan, 2003). For example, Tong and Flanagan (2003) reported interference at 5 min based on the average of the second and third cycles. Likewise, Miall and collaborators (2004) reported interference at 15 min based on the initial state obtained from fitting a power function, while noting that the rate of learning was not affected. Yet, there is evidence suggesting that using the initial level of performance as a proxy for anterograde interference may confound the actual ability of learning in *B* with the bias of a lingering memory of *A*. For example, Thoroughman and Shadmehr (1999) have shown that the preferred direction of the biceps and triceps during exposure to the second opposing force field is appropriate to solve the first force field. Moreover, Sing and Smith (2010) have demonstrated that the magnitude of initial errors does not impact on the ability to learn. In fact, high initial error may be associated either with faster (Sing et al., 2009) or slower learning. Therefore, assessing anterograde interference based on the initial level of performance may overestimate its magnitude. To rule out this possibility here we compared the bias imposed by the memory of *A* with the deficit observed in the rate of learning. We reasoned that if, as suggested by previous work, the initial level of performance reflects the level of anterograde interference then the two measures should behave similarly as a function of time. In contrast, we found that initial performance was profoundly hindered throughout the 24 h of testing, whereas the ability to learn returned to control levels by 6 h. This impairment in the ability to learn that gradually subsides with the passage of time is consistent with anterograde interference.

Our work sheds light on a long-standing debate regarding the failure of retrograde protocols at unveiling the time course of memory consolidation. Over the past two decades, several laboratories have attempted to uncover the time course of memory stabilization in sensorimotor adaptation using behavioral protocols based on retrograde interference (e.g. Brashers-Krug et al., 1996; Caithness et al., 2004; Krakauer et al., 2005). In these studies, subjects usually adapt to opposing perturbations *A* (*A*_**1**_) and *B* separated by a time interval that varies between minutes to 24 h. Next, they wait for a further period of time (usually 24 h) and are again exposed to *A* (*A*_**2**_) to assess the integrity of the motor memory. Consolidation of the memory of *A* should be reflected as the presence of savings (a faster rate of learning) in *A*_2_. Although this approach has proved successful in declarative (Lechner, Squire, & Byrne, 1999; Tulving, 1969) and some kinds of motor skill learning tasks (Korman et al., 2007; Walker, Brakefield, Hobson, & Stickgold, 2003), it has led to inconclusive results in sensorimotor adaptation. In fact, with the exception of three force-field studies reporting release from interference at around 6 h (Brashers-Krug et al., 1996; Shadmehr & Brashers-Krug, 1997) or later (Overduin, Richardson, Lane, Bizzi, & Press, 2006), other experiments have shown complete lack of savings even if 24 h are interposed between *A*_1_ and *B* (Caithness et al., 2004; Goedert & Willingham, 2002; Krakauer et al., 2005). Miall and collaborators (2004) have claimed that naïve performance at recall (*A*_2_) reported in retrograde protocols (Caithness et al., 2004; Goedert & Willingham, 2002; Krakauer et al., 2005) actually reflects a mixture of anterograde interference from *B* and the integrity of the memory of *A*, and not catastrophic retrograde interference. It is important to note, however, that these authors measured anterograde interference based on the initial level of performance. In light of our findings, the interpretation of these studies may need to be revised. Because release from interference occurs at 6 h anterograde interference is not likely to cause naïve performance in *A*_2_. Future work using retrograde protocols should in fact track the integrity of the memory based on the speed of learning in experimental designs in which *B* and *A*_2_ are separated by at least 6 h.

The temporal dissociation we observed between the initial level of performance and the rate of learning likely reflects two distinct processes at play: the persistence of a prior memory and competition for the same neural resources. The formation of memory involves learning-dependent synaptic plasticity as part of a process known as long-term potentiation (LTP). Given that biological substrates underlying synaptic plasticity are limited by nature, cellular modifications induced by learning temporarily constrain the capacity for further LTP induction. This phenomenon known as occlusion, which reflects competition from the same neural resources, has been implemented experimentally to investigate LTP maintenance and memory stabilization (Ling et al., 2002). Using this approach, it has been shown that motor skill learning in rats and humans is associated with LTP (Cantarero, Tang, O’Malley, Salas, & Celnik, 2013; Rioult-Pedotti, Friedman, Hess, & Donoghue, 1998; Rioult-Pedotti, Friedman, & Donoghue, 2000). Cantarero and collaborators showed that in fact, in humans, occlusion fades around 6 h after motor skill learning. In this light, we may speculate that adaptation in *A* may have partially occluded the capacity for further synaptic plasticity, thereby hindering adaptation in *B*. The timing of release from interference described herein (6 h) coincides with the peak in functional connectivity of a visuomotor adaptation network that includes the primary motor cortex (M1), the posterior parietal cortex (PPC) and the cerebellum (Della-Maggiore, Villalta, Kovacevic, & McIntosh, 2015). These regions have been linked to memory formation in this paradigm (Della-Maggiore, Malfait, Ostry, & Paus, 2004; Hadipour-Niktarash, Lee, Desmond, & Shadmehr, 2007; Landi, Baguear, & Della-Maggiore, 2011; Richardson et al., 2006). Whether this timing reflects the process of motor memory consolidation is now a hypothesis amenable for testing.

Using a state-space model allowed us to identify which aspect of learning was affected by anterograde interference. In other words, was the decrease in the rate of learning observed in *B* caused by a deficit in the ability to learn from error or in the ability to retain information cycle-by-cycle? Error sensitivity refers to how the brain responds to perceived error from one movement to the next and, as such, has been widely used to study changes across learning sessions in the context of savings and meta-learning (Herzfeld, Vaswani, Marko, & Shadmehr, 2014; Leow, de Rugy, Marinovic, Riek, & Carroll, 2016). The retention factor, on the other hand, refers to the degree of decay that occurs from one movement to another, reflecting the ability to retain information. It has been previously shown that the retention factor and error sensitivity can be independently affected by different manipulations (Galea, Mallia, Rothwell, & Diedrichsen, 2015) such as punishment (increasing only the sensitivity to error) and reward (enhancing retention). Here we found that prior learning of a contrasting perturbation had no effect on the cycle-by-cycle retention of subsequent learning.

Humans also have the capacity to change their error sensitivity depending on their prior experience with errors (Gonzalez Castro, Hadjiosif, Hemphill, & Smith, 2014). For example, Herzfeld and colleagues (2014) compared the sensitivity to error in participants that were exposed to alternating force fields, and found that individuals who experienced a slow switching environment showed a greater sensitivity to error than those who were exposed to a fast switching environment. These results indicate that the brain relies on a history of past errors to learn and, thus, when errors are not predictable learning is attenuated. Likewise, here we found that the temporal distance separating two opposing perturbations also hinders sensitivity to error. We may speculate that when perturbations are presented in close proximity (as in our 5 min and 1 h groups) competition for neural resources may hinder the ability to maintain a history of errors, resulting in a reduced error sensitivity. This phenomenon may revert at longer intervals as neural competition subsides.

In conclusion, we have examined the strength and duration of anterograde interference in visuomotor adaptation by tracking its impact on behavior when learning opposing perturbations separated from 5 min through 24 h. We found that prior learning dramatically hindered the initial state at all time intervals. This was likely due to a bias imposed by a lingering memory associated with the previous perturbation. Prior learning also impaired the ability to learn from error for at least 1 h, with release from interference detected about 6 h post training. The occurrence of release from interference within this time interval is consistent with a process of memory stabilization for this type of learning. Our findings suggest that poor performance observed when opposing rotations are learned subsequently is driven by two distinct phenomena operating on different time scales (days vs hours): a long-lasting influence of a memory that acts as a prior which negatively influences the initial level of performance, and a shorter-lasting impairment of learning.

## ACKNOWLEDGMENTS

This work was supported by the Argentinian Ministry of Defense (PIDDEF), the Argentinian Agency for the promotion of Science and Technology (FONCyT, ANPCyT), and the University of Buenos Aires (UBACyT). Reza Shadmehr was supported by grants from the National Institutes of Health (R01-NS078311, R01-NS095706). Scott Albert was supported by a pre-doctoral fellowship from the National Institutes of Health (NS-095706).

